# Memory Retrieval Effects as a Function of Differences in Phenomenal Experience

**DOI:** 10.1101/2023.07.31.551376

**Authors:** Austin H. Schmidt, C. Brock Kirwan

**Affiliations:** Neuroscience Center, Brigham Young University; Department of Psychology, Brigham Young University

**Author notes:** Correspondence should be addressed to Brock Kirwan.

**Keywords:** individual differences, phenomenal experience, experience, recognition, internal representations questionnaire, cognitive style

## Abstract

Conscious experience and perception are restricted to a single perspective. There is evidence to suggest differences in phenomenal experience can produce observable differences in behavior, however it is not well understood how these differences might influence memory. We used fMRI to scan n=49 participants while they encoded and performed a recognition memory test for faces and words. We calculated a cognitive bias score reflecting individual participants’ propensity toward either Visual Imagery or Internal Verbalization based on their responses to the Internal Representations Questionnaire (IRQ). We found weak positive correlations between memory performance for faces and a bias toward visual imagery and between memory performance for words and bias toward internal verbalization. There were typical patterns of activation differences between words and faces during both encoding and retrieval. There was no effect of internal representation bias on fMRI activation during encoding. At retrieval, however, a bias toward visualization was positively correlated with memory-related activation for both words and faces in inferior occipital gyri. Further, there was a crossover interaction in a network of brain regions such that visualization bias was associated with greater activation for words and verbalization bias was associated with greater activation for faces, consistent with increased effort for non-preferred stimulus retrieval. These findings suggest that individual differences in cognitive representations affect neural activation across different types of stimuli, potentially affecting memory retrieval performance.

---

Individuals differ in their phenomenological experience of the external world. This is also true for internal representations, some individuals expressing extremely vivid visual imagery (Cui et al., 2007; Marks, 1973) and others expressing an absence of visual imagery (i.e., aphantasia; Keogh & Pearson, 2018). Similarly, some individuals express strong internal verbalization (Alderson-Day et al., 2018) and others report an absence of internal verbalization (Heavey & Hurlburt, 2008) which has recently been termed “anendophasia” (Nedergaard & Lupyan, 2023). Recent evidence suggests that the phenomenological experience may influence performance in various cognitive domains. Participants who self-report having lower levels of inner speech have been shown to have lower performance on verbal working memory and rhyming judgment tasks (Nedergaard & Lupyan, 2023). Further, aphantasic participants, who report no ability to visualize despite no neurological damage, have been found to have generally reduced performance for episodic memory tasks, specifically retrieving episodic detail, compared to control groups (Blomkvist, 2023; Milton et al., 2021). Although previous literature has examined the effects of phenomenological experience for visual imagery and internal verbalization on memory, it is unclear how these experiences interact when forming and retrieving memories for verbal or visual information.

Prior research on individual differences in other domains has demonstrated effects on memory encoding and retrieval processes. For example, stable, non-strategic individual differences in recognition memory ability that are persistent over short (e.g., one week; Cohen, 1984) and long (e.g., 1-4 years; Woodhead & Baddeley, 1981; Zerr et al., 2018) delays. Further, individual differences in personality (Megreya & Bindemann, 2013) and mindfulness (Giannou et al., 2020) have been associated with memory performance for faces and encoding strategy (Karis et al., 1984) has been associated with memory performance for words.

A recently-developed tool called the Internal Representations Questionnaire (IRQ) provides a measure of individual differences in phenomenological experience, namely the propensity to use one cognitive style over another (Roebuck & Lupyan, 2020). The authors identify four factors of internal representation: Visual Imagery, Internal Verbalization, Orthographic Imagery, and Representational Manipulation. IRQ sub-scale scores for Visual Imagery and Internal Verbalization correlated with performance at visual and language-based tasks, respectively (Roebuck & Lupyan, 2020). This new measure provides the opportunity to use a more empirical examination of cognitive style, for both visual and verbal preference using the same tool, to determine if it has an observable impact on memory test performance.

Likewise, it also provides an avenue to examine whether corresponding differences in neural responses exist. We sought to answer whether individual differences as measured by the IRQ would influence memory recognition performance for visual and verbal stimuli, and if such differences would produce differential activation at encoding and retrieval. Specifically, we hypothesized that individuals who had more of a preference for internal verbalization would have better memory performance for verbal stimuli and those who had more of a preference for visual imagery would have better memory performance for visual stimuli (i.e., faces). Further, we hypothesized that brain regions associated with linguistic and visual processing would demonstrate differential activation corresponding to these behavioral effects.

## Methods

### Participants

A total of 120 participants were recruited from the university and nearby community. All participants self-reported free of neurological or psychiatric diagnoses and met safety inclusion criteria for MRI scanning (e.g., no history of metal injury to the eye). Participants were originally recruited for a study on the impacts of handedness on memory function (handedness analyses reported elsewhere). Consequently, 61 participants were excluded from the present analyses due left-handedness (Edinburgh Handedness Inventory Scores < 40). Further exclusions included seven participants due to excessive motion in the MRI scanner; two due to low response rates in the memory task (nonresponses >2 SD above the mean); and one due to being a notably strong outlier for cognitive bias score (−3.9 SD). The final data analyses were performed with n = 49 (25 male; 24 female, mean age = 23.1, range 18 to 48).

### Procedure

Prior to MRI scanning, participants completed the Internal Representations Questionnaire (IRQ), which is composed of four subscales: Visual Imagery, Internal Verbalization, Orthographic Imagery, and Representational Manipulation. We focused on the Visual Imagery and Internal Verbalization subscales given our research interest in whether the propensity to engage in visual imagery or internal verbalization affected memory for verbal or non-verbal stimuli.

While in the MRI scanner, participants performed encoding tasks for words then faces in separate blocks, a semantic fluency task (data used in a separate analysis; in preparation), and retrieval blocks for words then faces. We collected T1- and T2-weighted structural scans as well as a field map scan following the semantic fluency task prior to the retrieval task blocks, resulting in a delay between encoding and retrieval of approximately 15 minutes.

For the encoding task, participants rated as “pleasant” or “unpleasant” a series of 100 words and then 100 faces in separate scan runs (Guerin & Miller, 2009). Both face and word stimuli were presented for 2.5 seconds each followed by a fixation cross for 0.5-1.5 seconds, jittered (Amaro & Barker, 2006). Stimulus order was randomized for both encoding and retrieval. Face stimuli were selected from a database (Minear & Park, 2004) to have a broad distribution of demographics (e.g., age, sex, ethnicity). Word stimuli were selected from the MRC psycholinguistic database (Coltheart, 1981) to have high familiarity and high concreteness. For the retrieval task, participants performed a recognition memory task for 100 targets (faces or words) and 100 foils. Stimuli were presented one at a time for 2.5 seconds while participants made “old/new” recognition memory judgments. The inter-stimulus interval was again a fixation cross for 0.5-1.5 seconds, jittered. Retrieval tasks were broken into two scan runs of 100 trials each, totaling 200 trials for words, and 200 trials for faces. Responses were recorded using a four-button MR-compatible response cylinder (Current Designs Inc.; Philadelphia, PA). Stimuli were displayed using an MR-compatible LCD monitor placed at the head of the MRI scanner viewable using an adjustable mirror attached to the head coil.

### MRI Scanning

All MRI imaging was performed on a Siemens 3 Tesla TIM Trio scanner (Erlangen, Germany), using a 32-channel head coil. Each participant contributed a T1-weighted MP-RAGE structural scan (176 slices; TR = 1900 ms; TE = 4.92 ms; flip angle = 9°; field of view = 256 mm; slice thickness = 1 mm; voxel resolution = .97× .97 × 1.0 mm; 1 average) and echo-planar imaging (EPI) scans for each of the task blocks (72 interleaved slices; TR = 1800 ms; TE = 42 ms; flip angle 90°; field of view = 180mm; slice thickness = 1.8 mm; voxel resolution = 1.8 × 1.8 × 1.8 mm; Multi-Band factor = 4).

### Imaging Data Analysis

Unless otherwise noted, data were analyzed with AFNI (version AFNI_19.2.22) and SPSS (v.28). DICOM images were converted to NIfTI using *dcm2niix* (Li et al., 2016) and defaced. NifTI files were then uploaded to BrainLife.io (Avesani et al., 2019) and preprocessed using FreeSurfer and fMRIprep pipelines. Detailed descriptions of the fMRIprep pipeline autogenerated by the program are in Supplemental Materials. Additionally, functional data were blurred with a 4 mm FWHM Gaussian blur and scaled to have a mean value of 100 and range 0-200. Large motion events, defined as TRs with Euclidean norm (ENORM) of the temporal derivative of motion estimates greater than 0.3, were censored from the time series along with TRs immediately before and after.

Separate single subject regression models were created for face encoding, word encoding, face retrieval, and word retrieval tasks. Encoding task behavioral regressors coded for subsequent hits, subsequent misses, and trials of no interest including nonresponses. Retrieval task regressors coded for hits, misses, correct rejection (CRs), false alarms (FAs) and trials of no interest. For all tasks, events were modeled as a canonical hemodynamic response function convolved with a boxcar function of 2.5-second duration. All regression models included regressors for motion and polynomial regressors coding for run (retrieval tasks had two scan runs) and scanner drift. Resulting statistical maps of fit coefficients (β-coefficients) were entered into group-level analyses. Whole-brain analyses were corrected for multiple comparisons by first performing Monte Carlo simulations where smoothness of the data was estimated from the residuals of the single-subject regression analyses (Cox et al., 2017). Family-wise error was set to p < .05 with a voxel-wise threshold of p < .01 and a spatial extent threshold k > 88 for all analyses.

### Behavioral Data Analysis

We calculated discriminability (d’) scores for faces and words separately as z(hits)-z(false alarms). Given our focus on the relationship between visual imagery, internal verbalization, and memory processes, we first z-transformed raw visual imagery and internal verbalization subscale scores from the IRQ. Previous research has demonstrated that visual imagery and internal verbalization IRQ scores are positively correlated (Roebuck & Lupyan, 2020). Accordingly, in order to identify the differential influence of verbalization and visualization, we created a cognitive bias score by taking the difference between visual imagery and internal verbalization z-scores, i.e., z(visual imagery) – z(internal verbalization). Thus, positive scores indicate a greater propensity toward visual imagery and negative scores indicate a propensity toward internal verbalization. SPSS was used to obtain bivariate Pearson Correlations for face d’ by cognitive bias and word d’ by cognitive bias.

## Data and Code Availability

MRI data are available at https://openneuro.org/datasets/ds004589. Code used to present stimuli and perform all analyses is available at https://osf.io/tr78u/.

## Results

### Behavioral Results

Overall memory performance was good with mean (SD) d’ of 1.5 (0.4) for faces (t[48] = 26.40, p<.001, 95%CI [1.40, 1.63]) and 2.9 (0.7) for words (t[48]=28.09, p<.001, 95%CI [2.70, 3.11]). Memory performance for faces was found to be weakly positively correlated with cognitive bias score (r = .081, p = .582), while performance for words was found to be negatively correlated with the cognitive bias score (r = -.206, p=.156). The difference in correlations failed to reach significance (z=-1.547, p=.061). In other words, participants with a greater bias toward Visual Imagery over Internal Verbalization performed slightly better on face recognition while the reverse was true for word recognition performance (see Figure 1).

**Figure 1.**
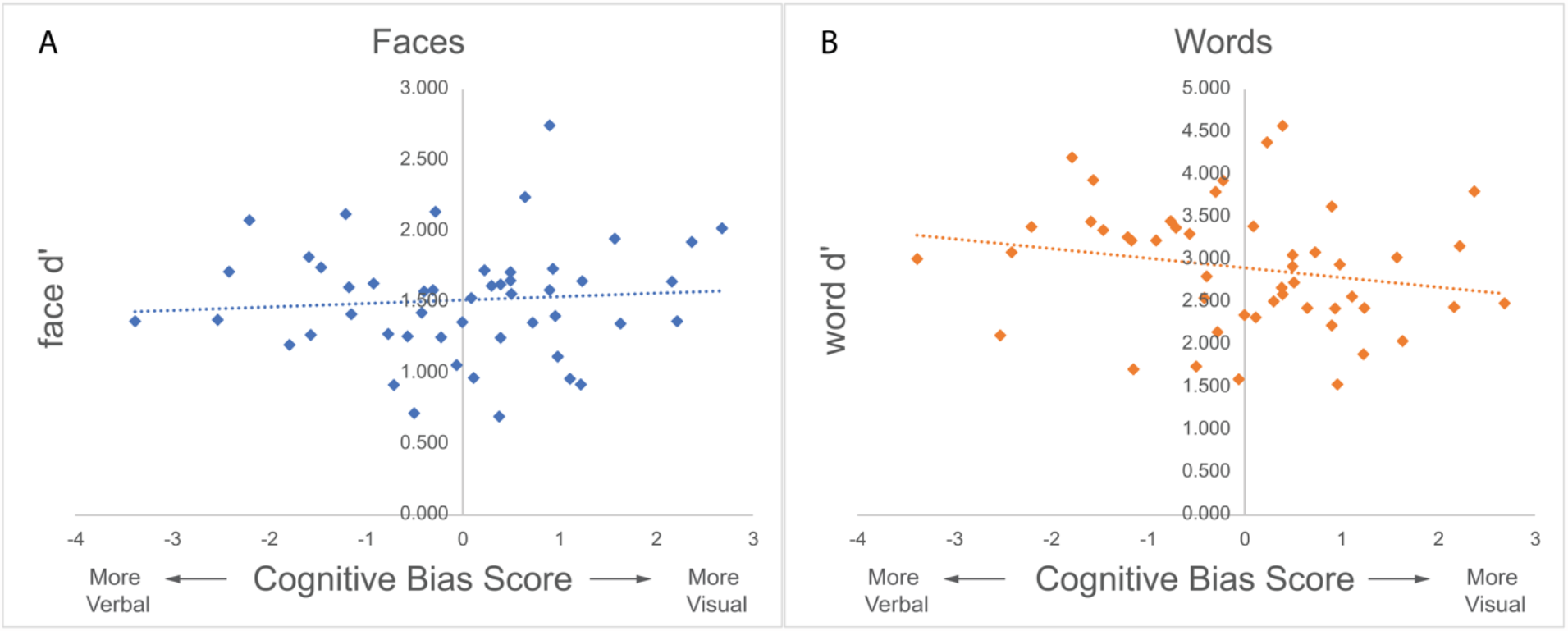
Memory Performance by Cognitive Bias Score. Scatter plots with regression line displaying discriminability (d’) scores for faces (A) and words (B) against individual cognitive bias scores. Positive scores indicate bias toward visual imagery. Greater visual imagery bias was associated with better face recognition memory performance and greater internal verbalization bias displayed the inverse association.

### Imaging Results

We performed separate analyses for encoding and retrieval tasks by performing whole-brain voxel-wise repeated-measures ANOVAs (AFNI program *3dMVM*) on memory-related activation with stimulus (words, faces) as a within-subject factor, and cognitive bias score as a continuous between-subjects factor. Memory-related activation at encoding was defined as subsequent hits minus subsequent misses. Similarly, memory-related activation at retrieval was defined as hits minus correct rejections. At both encoding and retrieval, there was widespread activation for the main effect of stimulus type, consistent with previous research demonstrating differential responding to faces compared to words (see Supplementary Materials). At encoding, there were no significant clusters for the main effect of cognitive bias or for the stimulus by cognitive bias interaction. At retrieval, there were two clusters of activation in bilateral inferior occipital lobe (left: 213 voxels, MNI coordinates -41, -91, -8; right: 97 voxels, MNI coordinates 33, -97, -7) that demonstrated a significant main effect of cognitive bias. In both clusters there was a positive correlation, such that a cognitive bias toward visualization was associated with greater retrieval activation, collapsing across faces and words (Figure 2). The correlation was significant in both left (r = .542, p < .001) and right (r = .480, p<.001).

**Figure 2.**
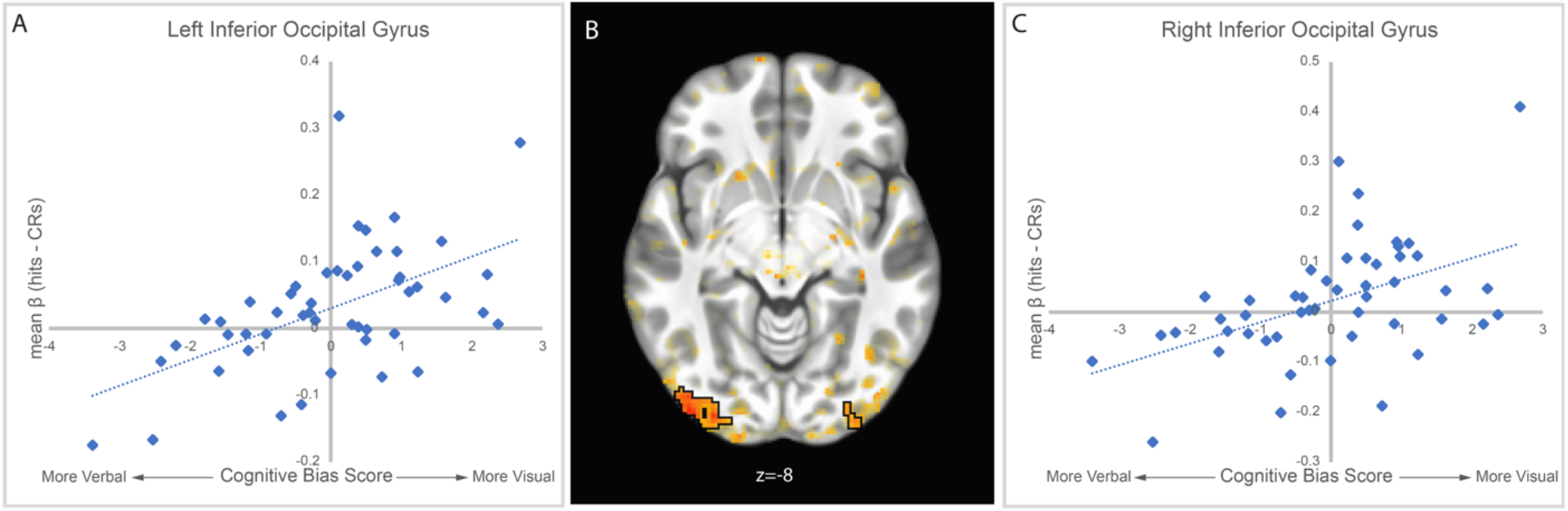
Main effect of cognitive bias at retrieval in left (A) and right (C) inferior occipital gyrus (B). Memory retrieval related fMRI activation was defined as Hits-CRs (collapsed across faces and words). A positive cognitive bias score reflects an individual’s propensity toward visual imagery. In both left and right inferior occipital gyrus there was a positive relationship between visual imagery bias and memory retrieval related fMRI activation.

Additionally, there were five clusters of activation at retrieval that demonstrated significant crossover interactions for cognitive bias by stimulus type (Table 1). These clusters included left lateral middle temporal gyrus, left inferior frontal gyrus orbitalis, left middle orbital gyrus, right inferior frontal gyrus orbitalis, and right angular gyrus (Figure 3A). These clusters all demonstrated a similar pattern of activity where a greater bias toward internal verbalization was associated with greater memory-related activation for faces, and a greater bias toward visual imagery was associated with greater memory-related activation for words (Figure 3B).

**Table 1.**

| Region | Number of Voxels | MNI Coordinates |  |  | Interaction |  |
| --- | --- | --- | --- | --- | --- | --- |
|  |  | X | Y | Z | F(1,47) | p |
| L. Lateral Middle Temporal Gyrus | 189 | -68 | -28 | -3 | 23.40 | <.001 |
| L. Inf. Frontal Gyrus Orbitalis | 172 | -41 | 31 | -5 | 26.32 | <.001 |
| L. Middle Orbital Gyrus | 135 | -44 | 55 | -1 | 20.46 | <.001 |
| R. Inf. Frontal Gyrus Orbitalis | 98 | 31 | 35 | -16 | 20.41 | <.001 |
| R. Angular Gyrus | 90 | 53 | -61 | 46 | 12.15 | .001 |

Note: L.=left; R.=right; Inf.=inferior

**Figure 3.**
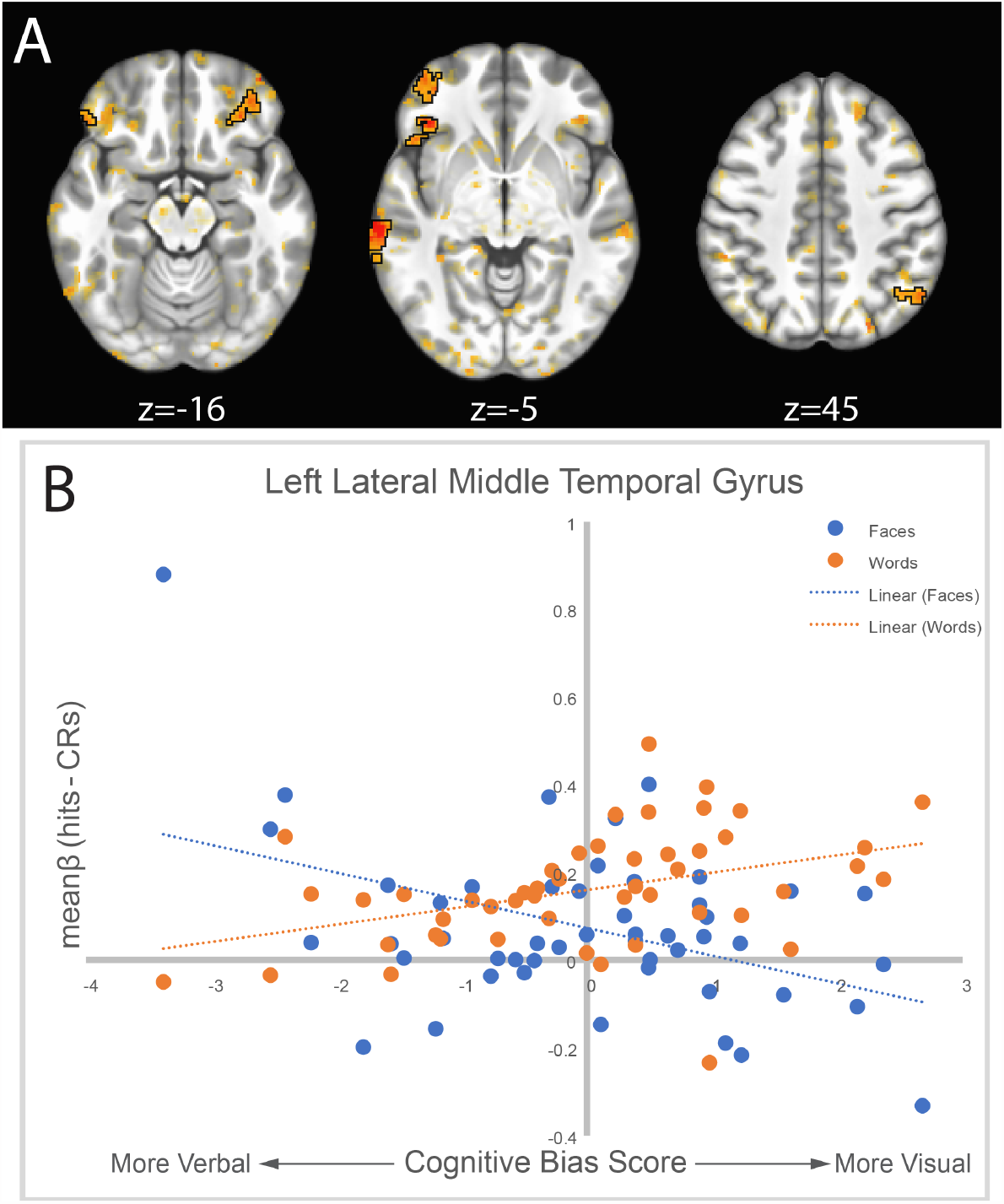
Clusters demonstrating a cognitive bias by stimulus type interaction at retrieval (A). All clusters followed a similar pattern of activity, illustrated by the activation from the largest cluster in left lateral middle temporal gyrus (B). Visual imagery bias was associated with greater activation for words, and less activation for faces. Internal verbalization bias was associated with the opposite pattern.

## Discussion

The primary aims of this study were twofold. First, we asked whether phenomenal experience as measured by the cognitive bias score would have a relationship with memory performance. Second, we asked whether the cognitive bias score would be associated with differential activation at encoding or retrieval for verbal and non-verbal stimuli. Our results for the former indicate a trend suggesting a relationship exists, but these findings are not significant. We did not observe any effects of phenomenal experience on fMRI activation for memory encoding regardless of stimulus type. On the other hand, during memory retrieval we found evidence for a main effect of cognitive bias in the inferior occipital gyri, such that higher visual bias was associated with greater memory retrieval related activation for both words and faces.

Additionally, we also observed a network of brain regions that displayed a cross-over interaction of cognitive bias by stimulus type, discussed below.

### Behavioral Findings

Our research question stems from previous findings demonstrating relationships between individual differences in non-mnemonic cognition and memory performance. Here, we asked if individual differences corresponding to phenomenal experience for verbal and visual information were associated with recognition memory performance for verbal and visual stimuli. Although we identified a general trend for memory performance in line with our hypothesis (Nedergaard & Lupyan, 2023; Woodhead & Baddeley, 1981), namely a higher verbalization bias associated with better word discriminability and higher visualization bias associated with better face discriminability, these results were not statistically significant. We suggest that the current study may not be sufficiently powered to answer the question of whether cognitive style affects memory performance (Anderson et al., 2017). The data that were utilized in this study derive from a larger project on the effects of handedness and when left-handed participants were included in the behavioral analysis the relationship was statistically significant (z = -2.15, p = 0.016). However, as handedness may affect fMRI activation patterns, we excluded these subjects from both behavioral and fMRI analyses to maintain consistency.

### Neural Findings

We observed greater activation (Hits-CRs) during memory retrieval in the inferior occipital gyri for visual biased participants collapsed across faces and words. This result may indicate that a propensity toward visual imagery produces greater recruitment of visual regions for memory retrieval tasks in general. This is consistent with previous research that has demonstrated a connection between visual imagery and face perception in occipital activation (Ishai, 2002; Slotnick et al., 2012). Greater activation in these areas for memory retrieval may suggest that individuals with a bias toward visual imagery are more likely to recruit occipital areas as they reconstruct a memory in a visual form regardless of the task format itself.

We also found a significant cognitive style by stimulus type cross-over interaction at retrieval reflecting greater memory activity when stimulus type did not match preferred cognitive bias. Active regions overlapped with the language network (left middle temporal gyrus and inferior frontal gyrus), the frontoparietal control network (inferior frontal and middle orbital gyrus), and the default mode network (right angular gyrus). A recent model of memory retrieval (Kim, 2020) suggests that the frontoparietal control network supports retrieval and decision effort while the medial temporal lobe (MTL) supports retrieval of memory representations and the DMN supports the subjective experience of remembering. While we did not observe differential activation of the MTL (consistent with activation across stimulus types and cognitive biases), we did observe differential activation consistent with this account in the FPCN and DMN. Also consistent with an interpretation of greater effort are studies that demonstrate increased activation in similar networks for linguistic task demands in (Klimovich-Gray et al., 2017).

We did not observe any effects of cognitive bias on memory encoding. Prior research on individual differences for mindfulness demonstrated effects at retrieval and not encoding, suggesting differences in decision-making strategy (Rosenstreich & Ruderman, 2016). Cognitive bias may similarly influence memory-guided decision-making strategy. Alternatively, the phenomenal experience of memory retrieval may vary according to cognitive bias, where the experience of autonoetic consciousness (Tulving, 2002) corresponds to preferred cognitive bias. Future research will be needed to characterize this possibility. Similarly, further research may wish to examine if internal imagery has an interaction with word concreteness for memory recognition (Taylor et al., 2019).

## Conclusions

Our findings demonstrate an effect of phenomenal style on memory retrieval activation. Participants with a bias toward visual imagery differentially activated visual cortices during memory retrieval regardless of stimulus type. We observed a further pattern of activation consistent with task demands, as activation was greater when cognitive bias and task type differed. Taken together, our findings suggest that differences in phenomenal experience are reflected in neural activation in retrieval tasks. This variability in retrieval activation should be taken into account in our theorizing about how brain networks interact in support of episodic memory retrieval.

