## Supplemental Materials for "Memory Retrieval Effects as a Function of Differences in Phenomenal Experience"

**Supplementary Materials**

*Main Effects of Stimulus*

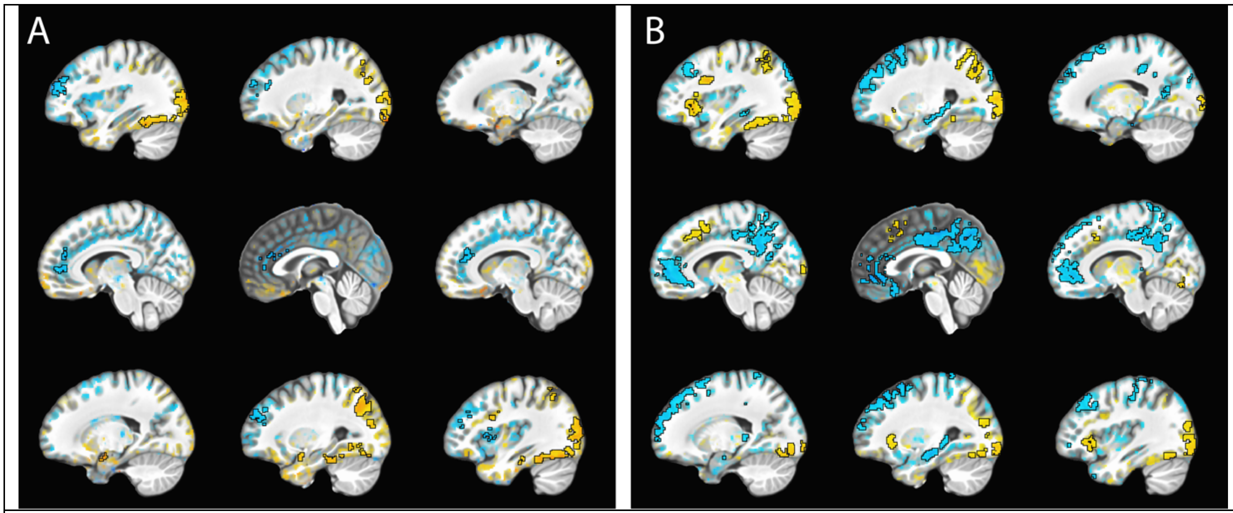


Figure S1. Main effect of stimulus type (defined as Faces > Words) during encoding (A) and retrieval (B).

*fMRIprep preprocessing pipeline*

Results included in this manuscript come from preprocessing performed using *FMRIPREP* version 20.2.1 [1, 2, RRID:SCR_016216], a Nipype [3, 4, RRID:SCR_002502] based tool. Each T1w (T1-weighted) volume was corrected for INU (intensity non-uniformity) using `N4BiasFieldCorrection` v2.1.0 [5] and skull-stripped using `antsBrainExtraction.sh` v2.1.0 (using the OASIS template). Brain surfaces were reconstructed using `recon-all` from FreeSurfer v6.0.1 [6, RRID:SCR_001847], and the brain mask estimated previously was refined with a custom variation of the method to reconcile ANTs-derived and FreeSurfer-derived segmentations of the cortical gray-matter of Mindboggle [21, RRID:SCR_002438]. Spatial normalization to the ICBM 152 Nonlinear Asymmetrical template version 2009c [7, RRID:SCR_008796] was performed through nonlinear registration with the `antsRegistration` tool of ANTs v2.1.0 [8, RRID:SCR_004757], using brain-extracted versions of both T1w volume and template. Brain tissue segmentation of cerebrospinal fluid (CSF), white-matter (WM) and gray-matter (GM) was performed on the brain-extracted T1w using `fast` [17] (FSL v5.0.9, RRID:SCR_002823).

Functional data was motion corrected using `mcflirt` (FSL v5.0.9 [9]). "Fieldmap-less" distortion correction was performed by co-registering the functional image to the same-subject T1w image with intensity inverted [13,14] constrained with an average fieldmap template [15], implemented with `antsRegistration` (ANTs). This was followed by co-registration to the corresponding T1w using boundary-based registration [16] with six degrees of freedom, using `bbregister` (FreeSurfer v6.0.1). Motion correcting transformations, field distortion correcting warp, BOLD-to-T1w transformation and T1w-to-template (MNI) warp were concatenated and applied in a single step using `antsApplyTransforms` (ANTs v2.1.0) using Lanczos interpolation.

Physiological noise regressors were extracted applying CompCor [18]. Principal components were estimated for the two CompCor variants: temporal (tCompCor) and anatomical (aCompCor). A mask to exclude signal with cortical origin was obtained by eroding the brain mask, ensuring it only contained subcortical structures. Six tCompCor components were then calculated including only the top 5% variable voxels within that subcortical mask. For aCompCor, six components were calculated within the intersection of the subcortical mask and the union of CSF and WM masks calculated in T1w space, after their projection to the native space of each functional run. Frame-wise displacement [19] was calculated for each functional run using the implementation of Nipype.

Many internal operations of FMRIPREP use Nilearn [22, RRID:SCR_001362], principally within the BOLD-processing workflow. For more details of the pipeline see https://fmriprep.readthedocs.io/en/20.2.1/workflows.html.

**Copyright Waiver**

The above boilerplate text was automatically generated by fMRIPrep with the express intention that users should copy and paste this text into their manuscripts *unchanged*. It is released under the [CC0](https://creativecommons.org/publicdomain/zero/1.0/) license.

4. Gorgolewski KJ, Esteban O, Ellis DG, Notter MP, Ziegler E, Johnson H, Hamalainen C, Yvernault B, Burns C, Manhães-Savio A, Jarecka D, Markiewicz CJ, Salo T, Clark D, Waskom M, Wong J, Modat M, Dewey BE, Clark MG, Dayan M, Loney F, Madison C, Gramfort A, Keshavan A, Berleant S, Pinsard B, Goncalves M, Clark D, Cipollini B, Varoquaux G, Wassermann D, Rokem A, Halchenko YO, Forbes J, Moloney B, Malone IB, Hanke M, Mordom D, Buchanan C, Pauli WM, Huntenburg JM, Horea C, Schwartz Y, Tungaraza R, Iqbal S, Kleesiek J, Sikka S, Frohlich C, Kent J, Perez-Guevara M, Watanabe A, Welch D, Cumba C, Ginsburg D, Eshaghi A, Kastman E, Bougacha S, Blair R, Acland B, Gillman A, Schaefer A, Nichols BN, Giavasis S, Erickson D, Correa C, Ghayoor A, Küttner R, Haselgrove C, Zhou D, Craddock RC, Haehn D, Lampe L, Millman J, Lai J, Renfro M, Liu S, Stadler J, Glatard T, Kahn AE, Kong X-Z, Triplett W, Park A, McDermottroe C, Hallquist M, Poldrack R, Perkins LN, Noel M, Gerhard S, Salvatore J, Mertz F, Broderick W, Inati S, Hinds O, Brett M, Durnez J, Tambini A, Rothmei S, Andberg SK, Cooper G, Marina A, Mattfeld A, Urchs S, Sharp P, Matsubara K, Geisler D, Cheung B, Floren A, Nickson T, Pannetier N, Weinstein A, Dubois M, Arias J, Tarbert C, Schlamp K, Jordan K, Liem F, Saase V, Harms R, Khanuja R, Podranski K, Flandin G, Papadopoulos Orfanos D, Schwabacher I, McNamee D, Falkiewicz M, Pellman J, Linkersdörfer J, Varada J, Pérez-García F, Davison A, Shachnev D, Ghosh S. Nipype: a flexible, lightweight and extensible neuroimaging data processing framework in Python. 2017. doi:10.5281/zenodo.581704.
